# Modelling Cell Shape in 3D Structured Environments: A Quantitative Comparison with Experiments

**DOI:** 10.1101/2023.08.07.552225

**Authors:** Rabea Link, Mona Jaggy, Martin Bastmeyer, Ulrich S. Schwarz

## Abstract

Cell shape plays a fundamental role in many biological processes, including migration, division and tissue morphogenesis, but it is not clear which shape model best predicts three-dimensional cell shape in structured environments. Here, we compare different modelling approaches with experimental data. 3D scaffolds for cell adhesion were manufactured using 3D laser nanoprinting and the shapes of single mesenchymal NIH/3T3 cells were compared by a Fourier method with surfaces that minimize area under the given adhesion and volume constraints. For the minimized surface model, we found marked differences to the experimentally observed cell shapes, which necessitated the use of more advanced shape models. We used different variants of the cellular Potts model, which effectively includes both surface and bulk contributions. The simulations revealed that the Hamiltonian with linear area energy outperformed the elastic area constraint in accurately modeling the 3D shapes of cells in structured environments. Explicit modeling the nucleus did not improve the accuracy of the simulated cell shapes. Overall, our work identifies effective methods for accurately modeling cellular shapes in complex environments, which in the future will enable the rational design of scaffolds for desired cell shapes.

**Author summary:** Cell shape and forces have emerged as important determinants of cell function and thus their prediction is essential to describe and control the behaviour of cells in complex environments. While there exist well-established models for the two-dimensional shape of cells on flat substrates, it is less clear how cell shape should be modeled in three dimensions. Different from the philosophy of the vertex model often used for epithelial sheets, we find that models based only on cortical tension as a constant geometrical surface tension are not sufficient to describe the shape of single cells in 3D. Therefore, we employ different variants of the cellular Potts model, where either a target area is prescribed by an elastic constraint or the area energy is described with a linear surface tension. By comparing the simulated shapes to experimental images of cells in 3D scaffolds manufactured with direct laser writing, we can identify parameters that accurately model the 3D cell shape.

## Introduction

The shape of animal cells is the result of active and passive intracellular and extracellular forces arising from actin polymerization, actomyosin contraction, cell adhesion, and material properties of the cell and its mechanical environment [1]. The cell membrane, a lipid bilayer with fluid-like properties, acts as the physical boundary of the cell and determines cell area and volume, but does not contribute directly to the mechanics of adherent cells. In most cell types, the actin cortex, a thin layer of actin filaments and myosin motor proteins located directly beneath the membrane, is the main mechanically relevant component [2]. The cortex actively contracts the cell surface, introducing tension gradients that enable the cell to change shape [3]. Other internal structures relevant to cell shape include actin stress fibers, which are contractile filament bundles that form dynamically in response to the mechanical environment, microtubules, stiff hollow structures that provide intracellular coordination and stability, and intermediate filaments, which contribute to cell integrity and resilience against external stress, mainly in epithelial cells [4–6]. Although our understanding of each system is increasing, the interplay between the three filament systems is crucial for cell mechanics and their resulting cell shape is difficult to predict due to the complexity of the system [7].

Cell shape is influenced by several factors. As different cell types ensure specific functions within the organism, their shapes are optimized for these functions depending on cell type. Furthermore, cell shapes change during development and morphogenesis. While all animal cells have the ability to change their shape in principle, cells that function alone, such as keratocytes and fibroblasts, are more dynamic than others and remodel their cytoskeleton on a minute time scale, enabling them to change their shape frequently. Additionally, cell shape changes are necessary during division [9]. Besides internal factors, the mechanical environment surrounding cells can also greatly influence their shape. With recent technological advancements, it is now easier to obtain 3D images of cells, enabling the investigation of not only their 2D projections, but also their full 3D shape [10].

Modeling plays an important role in increasing and validating our understanding of cell shape. Simulating cell shape in structured environments tests and increases our understanding of the underlying mechanisms governing cell behavior. Analytical 2D cell shape models have been developed to predict cell shape on micropatterned environments using line and surface tensions [17, 18], and the notion that the contractile actin cortex is responsible for 3D cell shape is wide-spread [19, 20]. In fact this philosophy underlies the popular vertex models for epithelial sheets and 3D cell assemblies, which reduces cell mechanics to the contractility of the cellular interfaces [21, 22]. 3D cell shape simulations have relied also on different approaches, including neural networks [23] and learned probability distributions [24]. In this study, we employ energy-based descriptions to model cell shape in well-defined environments and compare different surface energy descriptions. Our approach provides a robust framework for modeling complex 3D cell shapes and has the potential to improve our understanding of the fundamental principles that govern cell behavior in structured environments.

In order to quantify cellular shapes in structured environments, we utilized 3D structures manufactured with laser nanoprinting, which allowed us to create precisely controlled conditions [25]. First, we compare the experimentally observed shapes of single mesenchymal NIH/3T3 cells in structured environments with minimized surfaces under volume constraint using a Fourier decomposition. We found that these shapes can differ significantly, confirming that cell shape is more complex than surfaces under constant tension (surfaces of constant mean curvature (CMC)). To address this issue, we employed two different modeling approaches using the cellular Potts model (CPM) with elastic or linear area energies. Our simulations showed that the Hamiltonian with linear area energy outperformed the elastic area constraint in accurately modeling the shapes of cells in structured environments. We also found that explicitly modeling the nucleus did not necessarily improve the accuracy of the simulated cell shapes. Overall, our study provides insights into effective methods for modeling cellular shapes in complex environments.

## Materials and methods

### 3D Structured Environments

We manufactured structured environments using 3D laser nanoprinting [25]. In this technique, a photopolymerizable resist forms radicals in the focal volume of a femtosecond-pulsed laser. The resist only polymerizes after a two-photon absorption, and because the probability for a two-photon polymerization is only high enough in the focal volume of the laser, complex structures can be printed with micrometer precision.

The fabricated structures consisted of L-shaped, rectangular, V-shaped and triangular shaped patterns, see Fig. 1a-a*^′′′^*,b-b*^′′′^*, with 15 µm high anti-adhesive columns connected by biofunctionalized cross struts of 5 µm width, providing a suitable platform for cell adhesion in 3D. These geometries were selected to resemble the shapes of 2D micropatterns, which are well characterized in terms of 2D cell shapes [26], while eliminating the apico-basal polarity observed in cells on substrates.

**Fig 1.**
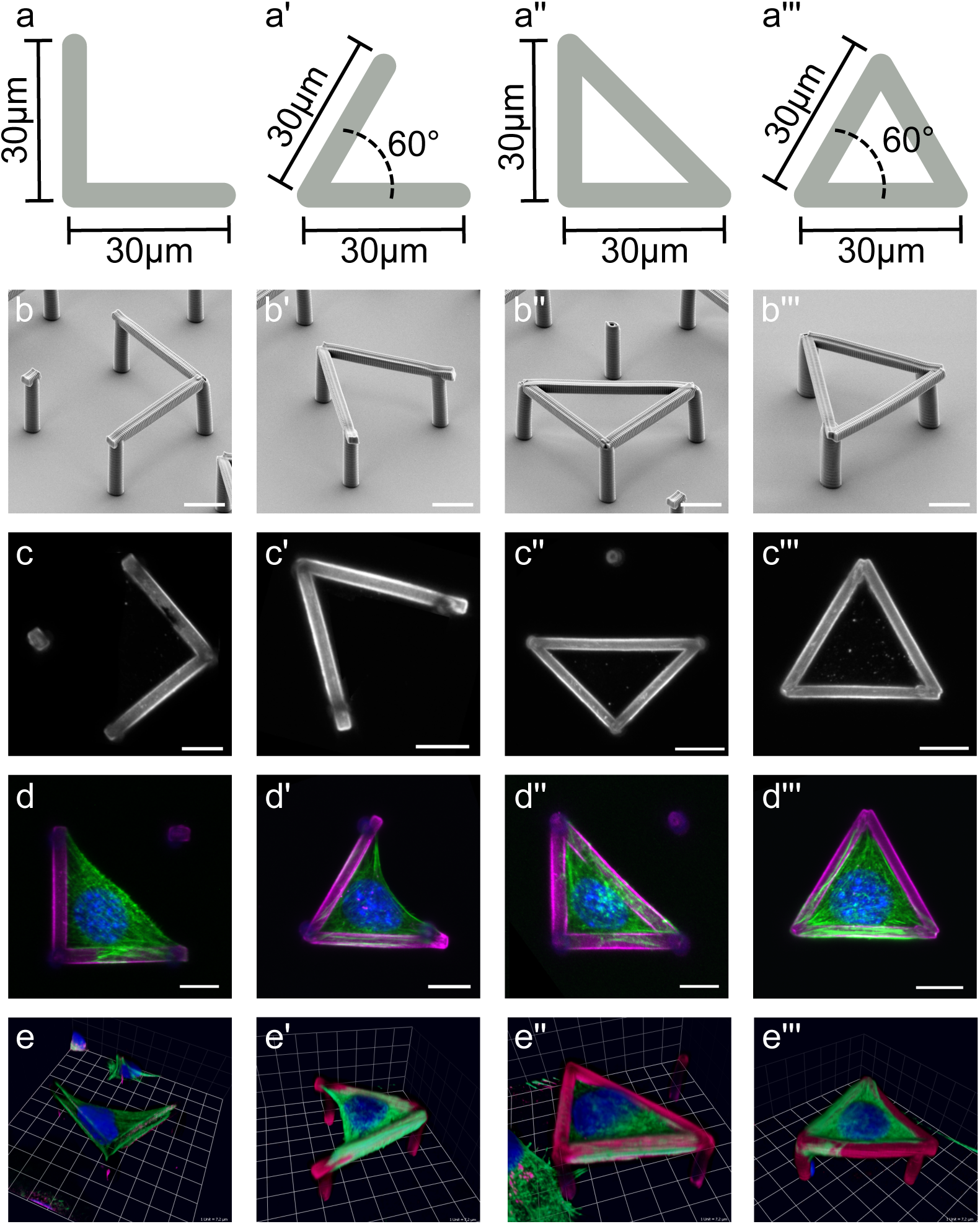
Overview of the structures used in this work. **a-a***^′′′^*) Top view with dimensions of the 3D structures. Shown in gray is the contact area (”cross struts”) for the cells in the L-shaped (**a**), V-shaped (**a***^′^*), right triangle-shaped (**a***^′′^*) and equilateral triangle-shaped (**a***^′′′^*) structures. The width of the struts is 5 µm. The supporting columns (not depicted) have a height of 15 µm. The angles not explicitly indicated are 90° and 45°, respectively. **b-b***^′′′^*) Scanning electron microscope images of the 3D structures. **c-c***^′′′^*) Fluorescence imaging of the structures after fibronectin coating and immunohistochemical staining. **d-d***^′′′^*) Fluoresence imaging of NIH/3T3 cells in fibronectin-coated structures after immunohistochemical staining (fibronectin = magenta, self-fluorescence of columns = blue, DAPI = blue, actin = green). **e-e***^′′′^*) 3D reconstruction of the cells in 3D structures. The cells adhere to the cross struts, using the entire strut as the adhesion surface. Scaffold not depicted in **e**. Scale bar: 10 µm.

To functionalize the structures, they were first rinsed with 70% ethanol (Carl Roth) and then dried for 30 min under UV light. Thereafter, the structures were overcoated with 200 µg*/*mL poly-L-lysine (Sigma-Aldrich) dissolved in phosphate-buffered saline (PBS, Biochrom AG) for 1 h at room temperature and then washed three times with PBS. This was followed by incubation with 10 µg*/*mL fibronectin in PBS for 1 h at room temperature. The functionalized structures can be seen in Fig. 1c-c*^′′′^*. After washing again three times with PBS, the structures were either used directly or stored in PBS for a maximum of two days at 4*^◦^*C.

### Cell Culture

NIH/3T3 embryonic mouse fibroblast cells were cultured at standard conditions (saturated humidity, 37*^◦^*C, 5% CO_2_) in serum containing medium and passaged three times per week to avoid contact inhibition. During passaging, cells were first rinsed twice with pre-warmed PBS and then incubated with 250 µL, 5% trypsin / 10 mM EDTA (Invitrogen) at 37*^◦^*C for 3 min to allow cells to detach from the substrate. The cell suspension was taken up in 5 mL of pre-warmed DMEM (Invitrogen) containing 10% fetal calf serum (FCS, PAA Laboratories). The serum in the medium provides saturation of trypsin. The cells were then centrifuged at 1000 rpm for 5 min. The supernatant was removed and the cell pellet resuspended in 5-10 mL of medium. Depending on the desired dilution, a certain volume was transferred to new cell culture flasks with pre-prepared tempered medium. The usual division ratios for NIH/3T3 cells is between 1:10 and 1:20.

After structure functionalization with fibronectin, NIH/3T3 mouse embryonal fibroblasts were seeded on the structures using a micromanipulator (aureka, aura optik GmbH) with an attached hydraulic manual microinjector (CellTram, Eppendorf). Cells were added to 4*^◦^*C CO_2_ buffered F12- imaging medium (0.76 g F12 nutrient mixture (Invitrogen), 50 mL water, 25mM HEPES (Carl Roth), 1% Pen/Strep (Sigma-Aldrich), 200mM L-glutamine (Life Technologies), 10% FCS). A portion of this cell suspension was pipetted over the glass plate containing the structures, which was also coated with F12 imaging medium (4*^◦^*C) and clamped in a magnetic holder. The unheated medium reduced premature adhesion of the cells to the substrate bottom and to the glass capillary of the microinjector. The individual cells were then aspirated to a glass capillary via the microinjector by creating a negative pressure. The capillary was then conveyed over the desired structure using the micromanipulator and the cell was transferred. After all structures were occupied, the temperature was raised to 37*^◦^*C to accelerate adhesion of the cells to the structures.

Protein staining was performed immunologically on fixed samples in a humidity chamber. Cells were fixed at room temperature for 10 min with 37*^◦^*C tempered 4% PFA (Sigma-Aldrich) in PBS. This was followed by permeabilization of the cell membrane by washing three times with 0.1% TritonX-100 (Carl Roth) in PBS for 5 min, followed by incubation with anti-fibronectin (BD Transduction Laboratories, 1:500) for 1 h at room temperature or at 4*^◦^*C. All antibodies and staining substances were diluted in 1% BSA (Bovine Serum Albumin) in PBS. This was followed by three wash steps for 5 min each with PBS. The fluorescent secondary antibodies anti mouse AF 647 (Life Technologies, 1:400) as well as DAPI (Roth, 1:2000) and fluorescently coupled phalloidin AF 488 (Life Technologies, 1:200) were then applied for 1 h at room temperature. After another wash step with PBS, the samples were embedded in 1% n-propyl gallate (Sigma-Aldrich) in Mowiol (Hoechst) and stored at 4*^◦^*C.

3D images of the cells were taken on the LSM 510 Meta (Zeiss) and the Axio Imager.Z1 with ApoTome (Zeiss) at 37*^◦^*C. The 3D shapes were extracted from the actin, DAPI and fibronectin staining as triangulated meshes using Imaris (Bitplane).

### Constant Mean Curvature (CMC) Surfaces

The actin cortex is a network of actin filaments and myosin motors positioned beneath the cell membrane, which contracts the cortex and influences cell shape. Assuming constant tension throughout the cortex, cell shapes can be characterized as fixed-volume objects that minimize their surface area, representing an approach that does not consider any contributions from the bulk of the cell. This approach is applicable to cells in suspension, which tend to be spherical in shape. For cells on adhesive stripes, their shapes can be described as a wetting process governed by surface tensions [27]. However, a quantitative comparison between experimentally observed cell shapes and shapes obtained from the described minimal surface model has not yet been conducted to our knowledge.

Surface minimization under volume constraints leads to surfaces with constant mean curvature (CMC) [28]. The mathematical problem of finding the minimum energy shape for a given boundary is called the Plateau problem. To assess the validity of these assumptions for cells in structured environments, we perform a comparison between experimentally observed cell surfaces and minimized surfaces with the same volume. The discrepancy between the observed and minimized cell shapes provides a measure of the degree to which factors beyond constant surface tension due to the actin cortex contribute to cell shape in structured environments.

The triangulated surfaces obtained from the experimental data were analyzed using SurfaceEvolver (Version 2.70) [29], a software that utilizes gradient descent to minimize surfaces under forces and constraints. Initially, the triangulated meshes obtained from Imaris were simplified using quadratic edge collapse decimation [30] and then used as inputs to SurfaceEvolver. The observed volume was fixed, as well as all points attached to the 3D printed scaffold. It was assumed that the surface tension is constant everywhere. The surfaces were minimized using the gradient descent method, and the minimized triangulated meshes were exported for further analysis.

### Cellular Potts Model (CPM)

Cellular Potts models (CPMs) are a more evolved approach to model single cells or cell collectives. Introduced by Glazier and Graner in the 1990s to model differential adhesion [31], CPMs are predominantly used to describe phenomena arising due to cellular interaction [32–34], but they have also been used to describe single cell migration and dynamics on micropatterns [18, 35–37]. In contrast to the surface model, they include bulk terms representing also the volume elasticity of cells.

In detail, cells are modeled on a lattice where each lattice site can be occupied by a generalized cell *σ* of a predefined cell type *τ*, which represents biological cells or other entities such as mechanical structures, or the medium (*σ* = 0). A cell typically occupies many lattice sites. The Hamiltonian ℋ is an energy functional that defines the energy for all possible lattice configurations. During the simulation, a modified Metropolis algorithm is used to minimize the total energy of the system. The algorithm attempts to update the configuration of the system by selecting a lattice site and trying to change its state to that of a neighboring generalized cell. The change is accepted with a probability given by the Metropolis rule, which depends on the energy difference between the new and the old configurations. By repeating this process many times, the system evolves to its lowest energy state [38].

Choosing the appropriate Hamiltonian to describe biological systems has been a longstanding question in the field. Early formulations of the Hamiltonian included an elastic volume constraint and interaction energies. In order to more accurately model cell behavior, an elastic surface constraint was added. Additionally, the nucleus can be modeled explicitly as a compartmentalized cell with an elastic constraint to ensure that the nucleus is close to the cell center of mass:

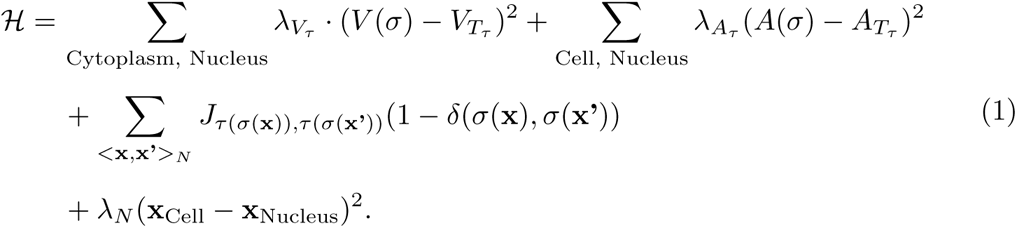

Here, the first two terms are the elastic volume and surface energies for the generalized cell types “cytoplasm” and “nucleus”, where the strength of the constraints is given by *λ_Vτ_* and *λ_Aτ_* respectively and the target volume and area are given by *V_Tτ_* and *A_Tτ_*. These parameters depend on the cell type *τ*. We describe the cytoplasm and the nucleus as generalized cell types each, and the scaffold is implemented as a fixed generalized cell. The inclusion of an elastic volume constraint in the Hamiltonian is motivated by the constant volumes of cell compartments in biological systems, and the need to model dynamics with some flexibility to avoid “lattice freezing” while ensuring that the simulated cells do not disappear, which would reduce the energy of the system but does not align with biological reality. Conversely, the elastic area constraint is not strictly required for the simulation, but its inclusion provides an additional constraint that allows more flexibility in the parameter selection of the interaction energies *J* [38]. The third term in the Hamiltonian represents the interaction energy at cell interfaces. It is computed by summing over all voxels **x, x’** within the neighborhood *N*, which for *N* = 1 are the 6 directly adjacent voxels, for *N* = 2 it includes the 8 diagonally adjacent voxels and so on. If the voxels **x, x’** belong to different generalized cells *σ*, their interaction energy *J* is added to the total energy of the system. The value of *J* can be positive or negative, and depends on the cell types of the neighboring voxels. To ensure that the nucleus remains inside the cell, an elastic constraint with strength *λ_N_* is added between the cell center of mass **x**_Cell_ and the center of mass of the nucleus **x**_Nucleus_.

Instead of the elastic area constraint that follows from the assumption that the cell surface is constant throughout the experiment, one can describe the surface with a linear area term, which follows from the assumption that increasing the cell surface always costs energy [33, 36]:

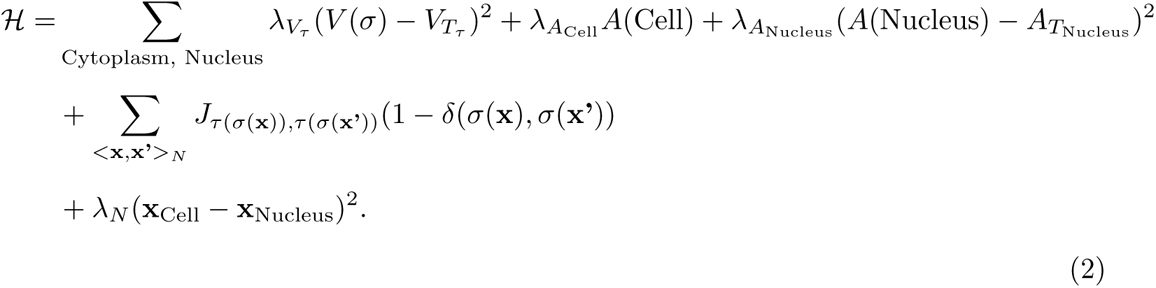

Here, the volume, interaction and nucleus surface and centering energy terms are similar to Eq. 1, but the elastic area energy term of the cell is replaced with a linear area energy, where the strength is determined by *λ_A_*_Cell_. Note that here, *λ_A_*_Cell_ has different dimensions compared to Eq. 1.

The simulations were implemented using CompuCell3D (version 4.1.1) [39]. The static cell shapes used for further analysis are obtained by averaging over the last 500 Monte Carlo Steps of the 2000 Monte Carlo steps simulation.

### 3D Spherical Harmonics Analysis

To analyze and quantitatively compare 3D cell shapes obtained from experiments and simulations, we employed a 3D spherical harmonics analysis. Spherical harmonics form a complete set of orthonormal functions and can be used as an orthonormal basis for describing 3D shapes. This approach enables precise, translation-invariant, scale-invariant, and rotation-invariant description of 3D shapes and has been previously used to analyze biological shapes [40–44].

Following the approach in [43], we first convert the boundary shapes for simulated and experimentally observed cell shapes from Cartesian coordinates to spherical coordinates, exploiting the fact that all our shapes are star-shaped and thus can be mapped bijectively to the unit sphere. We then sample the data on a regular grid and use the Driscoll and Healy sampling theorem [45] implemented in pyshtools (version 4.10) [46] to calculate the spherical harmonics:

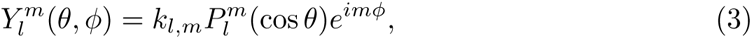

where *l* and *m* are the degree and order respectively. *k_l,m_* is the normalization and *P^m^* are the associated Legendre polynomials. The spherical harmonics as defined in Eq. 3 define an orthonormal basis, thus any scalar function *f* (*θ, ϕ*) on a sphere can be expressed as a sum of the spherical harmonics:

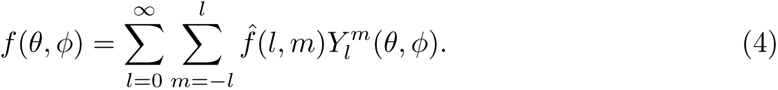

Here, 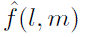 are the harmonic coefficients given by

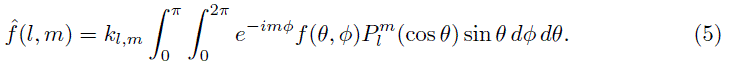

We normalize the harmonic coefficients 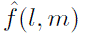 with respect to the first-order ellipsoid 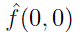 and use the normalized coefficients 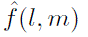 to calculate the rotation-invariant frequency spectrum *F* (*l*) as a quantitative shape measure

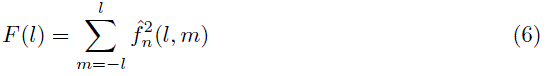

Now, we can calculate a measure for the difference Δ*_l_*_max_ between two shapes *a* and *b* from the corresponding frequency spectra:

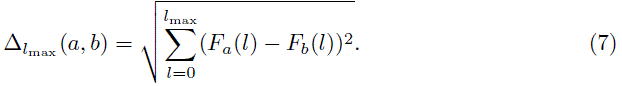

In the subsequent analysis, we will use the first 30 degrees of the respective frequency spectra to compare shapes (Δ_30_).

## Results

### Characterization of experimentally observed shapes

We first analyzed the shapes of *n* = 6 cells in L-shaped or square-shaped structures spanning between two fibronectin coated cross struts. A representative cell can be seen in Fig. 1d,e, and S1 Video. Visual inspection of the extracted cell shapes revealed remarkable similarity across all cells. Notably, the cells bridged along the free area between the beams, forming invaginated arcs – a well-established phenomenon observed for cells on both 2D microstructured surfaces and 3D structures [47, 48]. In some cases, we observe w-shaped invaginations due to the nucleus. Similarly, the shapes of *n* = 7 cells in V-shaped structures were analyzed, see Fig. 1d*^′^*,e*^′^* and S2 Video, where we again observe invaginated arcs. In this case, the nucleus does not interfere with the formation of the arc, and we do not observe w-shaped arcs. Additionally, we analyzed the shapes of *n* = 3 cells in right triangle structures, see Fig. 1d*^′′^*,e*^′′^*and S3 Video, and of *n* = 4 cells in equilateral triangle structures, see Fig. 1d*^′′′^*,e*^′′′^* and S4 Video. In these structures, the cells also adhere to the additional cross strut as compared to the L-shaped structure or the V-shaped structure respectively. In these cases, we do not find invaginated arcs as the cells fill the volume between the struts.

### Comparison to CMC-surfaces

We employed Imaris to extract triangulated meshes of the experimentally observed shapes, and combined surface of cytoplasm and nucleus for the purpose of surface minimization, see Fig. 2a-a*^′′′^*. To obtain the minimized surfaces, we used the triangulated mesh of the experimentally obtained cell surfaces and fixed vertices close to the structures. Then, we used SurfaceEvolver to minimize the shapes under a constant volume constraint. Upon visual inspection, we found significant differences between the experimentally observed cell shapes and the resulting minimal energy shapes for the L- and V-shaped structures, compare Fig. 2a,b and Fig. 2a*^′^*,b*^′^*, indicating that minimizing area under a volume constraint is insufficient for accurately describing cell shape in these cases. The observed cells in the scaffolds stretch between adhesive areas and maintain a roughly constant thickness, while the minimal energy surfaces form more sphere-like shapes with two thinly stretched extensions attached to the scaffold due to the imposed boundary conditions. This finding is unsurprising since spheres have the lowest surface-to-volume ratio, thus the surfaces are becoming sphere-like to minimize their surface area. For cells in the L-shaped and square scaffolds, this results in a w-type invagination. On the other hand, for cells in triangular shaped structures adhering to all three cross struts, the difference between the experimentally obtained and minimized surfaces are smaller, compare Fig. 2a*^′′^*,b*^′′^* and Fig. 2a*^′′′^*,b*^′′′^*. The surfaces are smoother, but cell-scale surface changes are not found.

**Fig 2.**
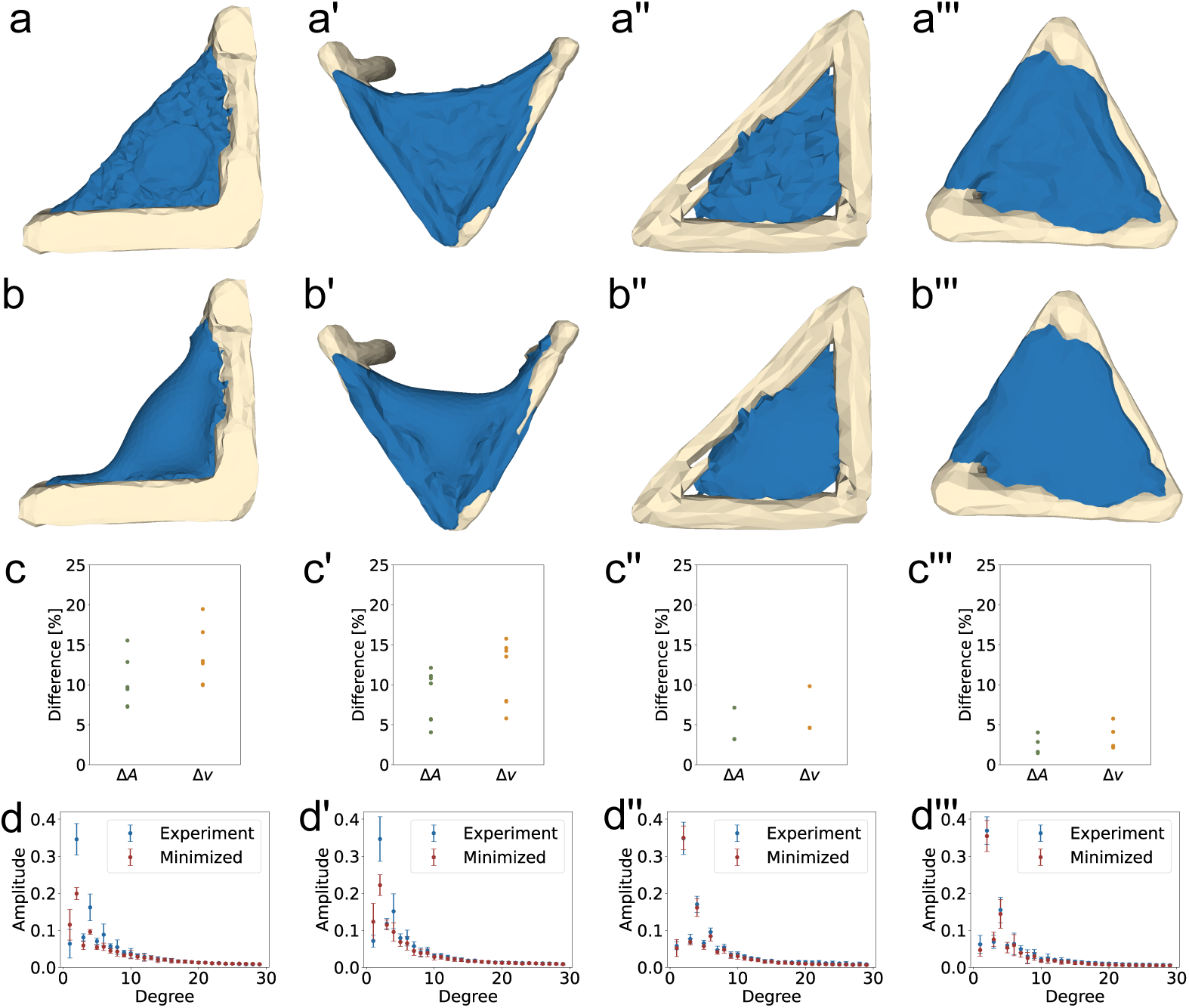
Comparison between experimentally observed cell shapes and their minimized surfaces. **a-a***^′′′^*) The surfaces of the cells in structured environments. To minimize the cell surface, the cytoplasm and nucleus surface reconstruction were combined to the cell surface. **b-b***^′′′^*) Minimized surfaces in the 3D structures. Mesh points attached to the structure were fixed during minimization. **c-c***^′′′^*) Area (Δ*A*) and reduced volume (Δ*v*) differences between the experimentally observed and minimized surfaces. **d-d***^′′′^*) Frequency spectrum of the experimentally observed (blue) and minimized (red) surfaces. **d)** Δ_30_ = 0.175. **d***^′^*) Δ_30_ = 0.148. **d***^′′^*) Δ_30_ = 0.026. **d***^′′′^*) Δ_30_ = 0.031. Movies of the 3D reconstructions and the minimized shapes can be found in S1 Video-S4 Video.

Landmark values of the surfaces before and after minimization quantify the shape differences. We compare the surface area reduction Δ*A* and the changes in reduced volume Δ*v*. The reduced volume *v*, which quantifies how much a shape differs from a perfect sphere, was calculated from 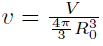, where *V* is the volume of the cell and *R*_0_ is computed from the surface area *A* of the cell as 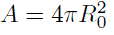. A reduced volume of 1 indicates a perfect sphere, while all other shapes have a lower reduced volume. The differences in area and reduced volume are illustrated in Fig. 2c-c*^′′′^*. For cells in the L-shaped structures, the surface area was reduced by (10*±*3)% during the minimization process on average, for the V-shaped structures the reduction was (9*±*3) %. Much smaller reductions were found for cells in triangular structures, cell area in the right angle triangle was reduced on average by (5*±*2) % and in equilateral triangles the difference is (2*±*1) %.

For the reduced volume *v*, we find similar differences. After minimization, the reduced volume of cells in L-shaped structures differed by an average of (14*±*3) % when compared to the reduced volume before minimization. For cells in V-shaped structures, this difference was (11*±*4) %, whereas for cells in right angle triangles, the average difference of Δ*v* was found to be (6*±*2) % and for cells in equilateral triangles (4*±*1) %.

These significant variations for cells that form invaginated arcs underscore the notion that 3D cell shape in structured environments is more complex than a surface that has been minimized under a volume constraint due to an isotropically contracting actin cortex.

To quantify these differences further, we subsequently computed the spherical harmonics coefficients of the obtained experimental and minimized shapes. To ensure that small surface fluctuations are not included in the analysis, a cutoff was applied after *l*_max_ = 30, see Eq. 7.

The frequency spectrum of the spherical harmonics for experimentally observed and minimized shapes can be seen in Fig. 2d-d*^′′′^*. The dipole moment of the spherical harmonics, represented by the amplitude of the first degree in the frequency spectrum, exhibits a low value. This result is anticipated, given the observed cell shape’s lack of symmetry with respect to a single axis. On the other hand, the second degree of the frequency spectrum displays the highest amplitude across all observed cell shapes, corresponding to the quadrupole moment that characterizes a distribution with two perpendicular axes of symmetry. Additionally, the fourth spherical harmonic also has a pronounced peak in the frequency spectrum, representing a six-axis symmetric distribution.

With the spherical harmonics analysis of the minimized structures, we are able to quantify the difference between experimentally observed shapes and what we would expect from minimal surfaces. The difference between the average experimentally observed and minimized amplitude is Δ_30_ = 0.175 for cells in L-shaped structures. For cells in V-shaped structures, we find Δ_30_ = 0.148. In both cases, there is a large difference in the low degrees of the frequency spectrum. The amplitude of the first degree of the frequency spectrum for the minimized shapes is larger than the experimental amplitude, while the amplitudes of the second and forth degrees, which are prominent in the experimental shapes, are lower. This confirms the increased sphericity of the minimized shapes. When comparing the experimentally observed cell shapes to the minimized shapes in triangular structures, the difference is much smaller; for cells in the right angle triangles, we find Δ_30_ = 0.026, and for cells in equilateral triangles we find Δ_30_ = 0.031. Both values are much lower than for structures with invaginated arcs, highlighting that the accuracy of cell shape prediction using constant mean curvature depends on the structured environment.

With this we show that 3D cell shape in structured environments is not always adequately described by minimized surfaces. We conclude that a constant surface tension due to a contractile actin cortex is not enough to describe cell shape, especially for cells with invaginated arcs.

### Comparison to CPM-shapes

The CPM can be used to simulate the behavior of cells in complex geometries, such as the scaffolds used in the experiments described above. By optimizing the parameters of the model it is possible to simulate cell shapes that closely resemble those observed experimentally. Parameters influencing the cell shape are the surface energy strength *λ_Aτ_*, which represents the energy required to change the cell surface area, as well as the interaction energy between the cytoplasm and the medium *J*_cytoplasm,medium_, which represents the energy required to move a cell in the surrounding medium. The neighbor order *N* is a parameter that determines the extent of the interactions between neighboring voxels. Choosing an appropriate neighbor order is crucial in CPM simulations, as it defines how much of the surrounding affects the cell, and to ensure that the neighborhood remains smaller than the cell [49]. Other parameters do not change the cell shape as long as their value is chosen within a reasonable range. The volume parameters *λ_Vτ_* and *V_Tτ_* are used to ensure that the volume of the cytoplasm and the nucleus stay close to the target volume *V_Tτ_* throughout the simulation, however changing the constraint strength *λ_Vτ_* within an appropriate range does not change the cell shape. The interaction parameters between cytoplasm and scaffold *J*_cytoplasm, scaffold_, as well as cytoplasm and nucleus *J*_cytoplasm, nucleus_, are chosen to ensure the cytoplasm adheres to the scaffold and the nucleus is surrounded by cytoplasm. The simulation temperature is fixed at *T* = 100 throughout the simulations. All parameters used in the simulations can be found in Table.

First, we investigate the effect of the neighbor order on cell shape. In Fig. 3, the measure of shape difference Δ_30_ between the experimentally observed cell shapes and the cell shapes simulated with the elastic area Hamiltonian (Eq. 1) for varying neighbor order *N* and interaction energy *J*_medium,cytoplasm_ can be seen. Remarkably, for a large parameter range of *J*_medium,cytoplasm_ and intermediate neighbor order *N*, significantly lower values of Δ_30_ than for the constant mean curvature shapes are found.

**Fig 3.**
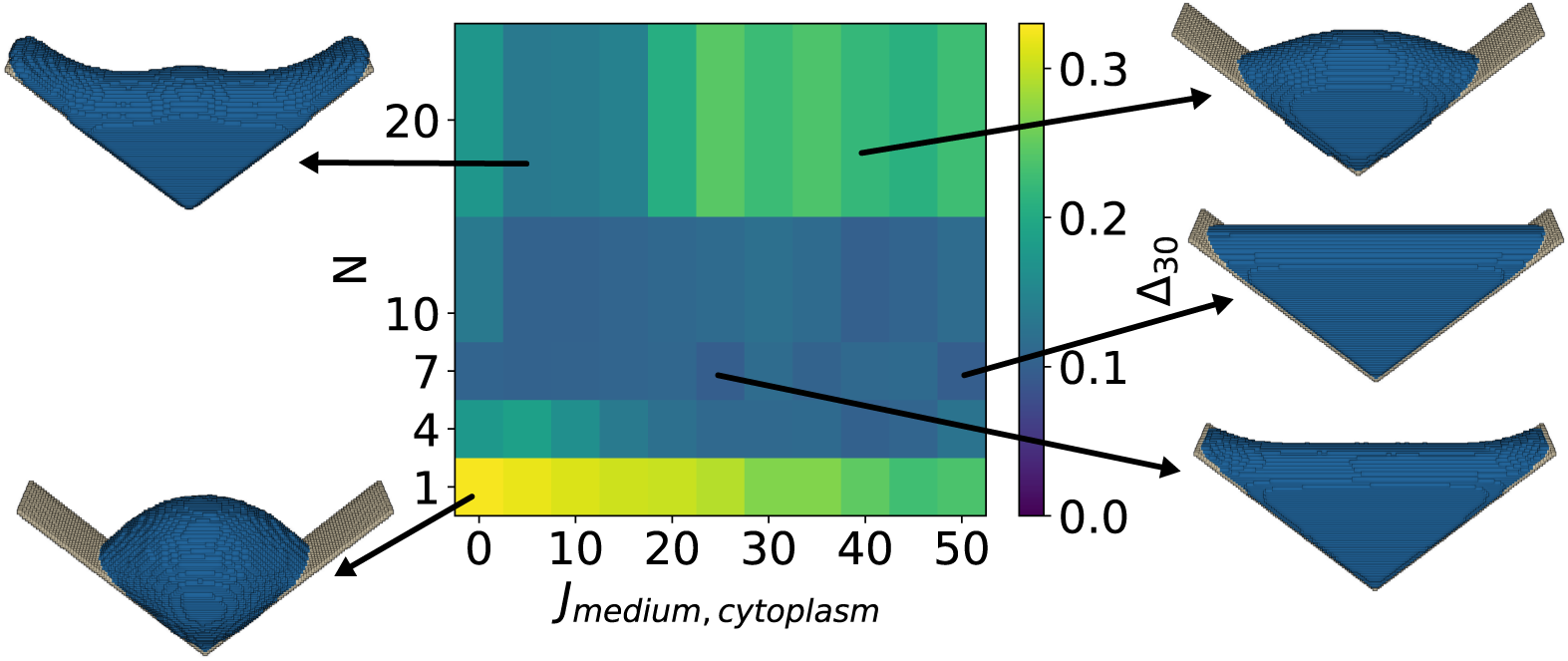
Cell shape difference Δ_30_ (Eq. 7) as a function of neighbor order and interaction energy. Neighbor order *N* defines the set of neighboring lattice sites that interact with a given lattice site, and interaction energy *J*_medium,cytoplasm_ defines the interaction strength between the cytoplasm and the surrounding medium. Simulated cell shapes for exemplary parameter choices are presented. The minimum cell shape difference Δ_30_ = 0.097 is found for *N* = 7 and *J*_medium,cytoplasm_ = 50, the corresponding cell shape is depicted on the bottom right.

The neighbor order parameter effects the simulated cell shape more than the interaction energy *J*_medium,cytoplasm_. When only the nearest neighbors are considered in the calculation of the interaction energy (*N* = 1), then the cell partially detaches from the scaffold due to a reduced energy gain from cytoplasm-scaffold interactions. Additionally, both the cytoplasm and nucleus become more spherical. For intermediate neighbor orders, 4 *≤ N ≤* 10, the simulated cell shapes resemble the experimentally observed shapes for a large parameter range of the interaction energy *J*_medium,cytoplasm_, leading to low values of Δ_30_, with the minimum being at Δ_30_ = 0.097, for *J*_medium,cytoplasm_ = 50 and *N* = 7. Increasing the neighbor order further to *N* = 20 results to shapes that deviate more from the experimentally observed cell shapes. When the interaction energy *J*_medium,cytoplasm_ is low, the shape becomes w-shaped and resembles that of minimized surfaces because the energy gain from cytoplasm-scaffold interactions is large due to the high neighbor order. At the same time, the remaining cytoplasm volume reduces its energy by becoming spherical, leading to a w-shape. However, when the interaction energy *J*_medium,cytoplasm_ is sufficiently increased, the cost of a less spherical shape becomes higher than the gain from the cytoplasm-scaffold interaction, resulting in more spherical shapes and increasing the value of Δ_30_ again. As the minimum of Δ_30_ is reached for *N* = 7, which is a neighbor order with a low perimeter scaling error due to the underlying lattice *J*_medium,cytoplasm_. Therefore, we fix the neighbor order to *N* = 7 in the following simulations.

One of the longstanding questions in CPM-type simulations is the selection of the appropriate Hamiltonian. Specifically, when it comes to surface area, the two commonly used Hamiltonians are either based on the assumption that the surface of a cell is approximately constant, thus describing the area energy with an elastic constraint, see Eq. 1, or that increasing surface area always costs energy for the cell, leading to a linear area energy, see Eq. 2. Comparing simulated cells with experimentally observed cell shapes allows for the direct comparison between the two approaches. Cell shapes simulated with either Hamiltonian, shown in Fig.4 for elastic area energy and in Fig.5 for linear area energy, closely resemble the experimentally observed cell shapes for a wide range of parameter values, vastly outperforming the minimized surface cell shapes.

**Fig 4.**
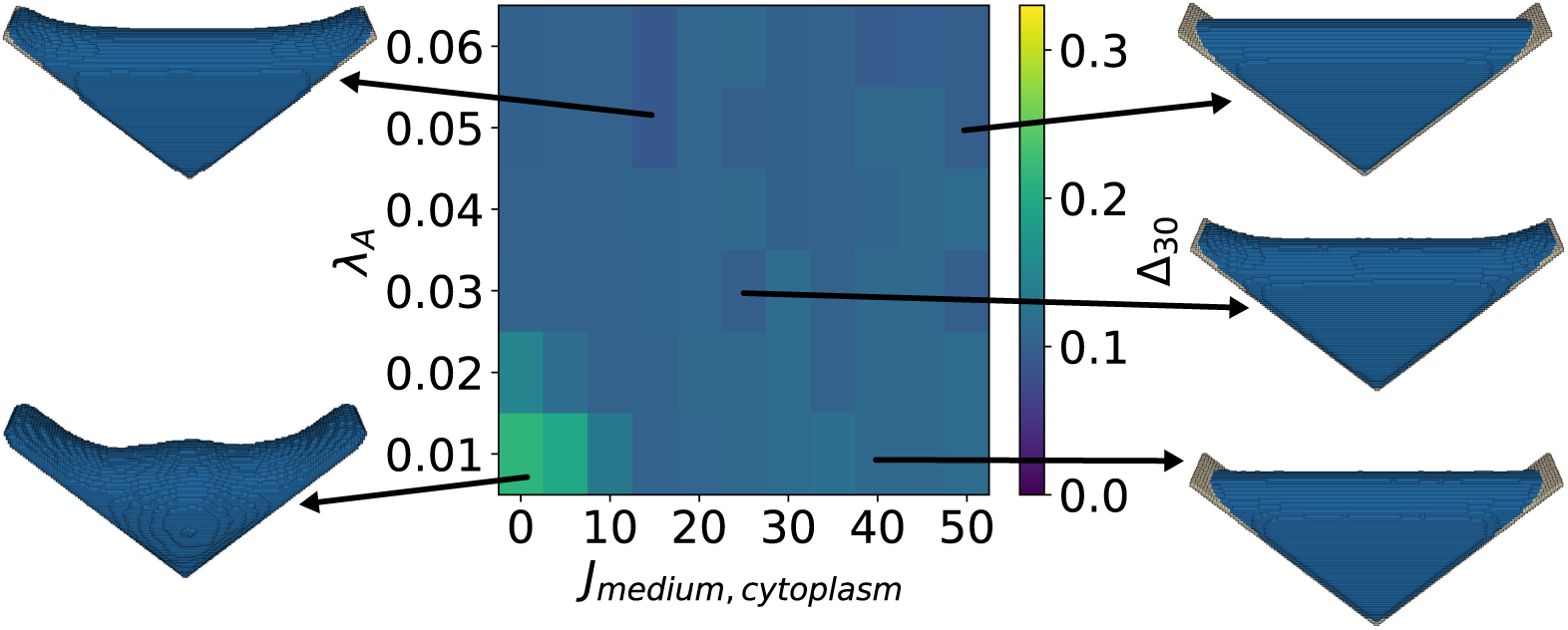
Cell shape difference Δ_30_ between the experimental and simulated cell shapes obtained with the elastic area energy (Eq. 1). Differences are shown as a function of surface energy constraint *λ_A_*and interaction energy *J*_medium,cytoplasm_. Simulated cell shapes for exemplary parameter choices are presented. The minimum cell shape difference Δ_30_ = 0.090 is found for *J*_medium,cytoplasm_ = 15 and *λ_A_* = 0.05, the corresponding simulated cell is depicted on the top left. A movie of the simulation can be seen in S5 Video.

**Fig 5.**
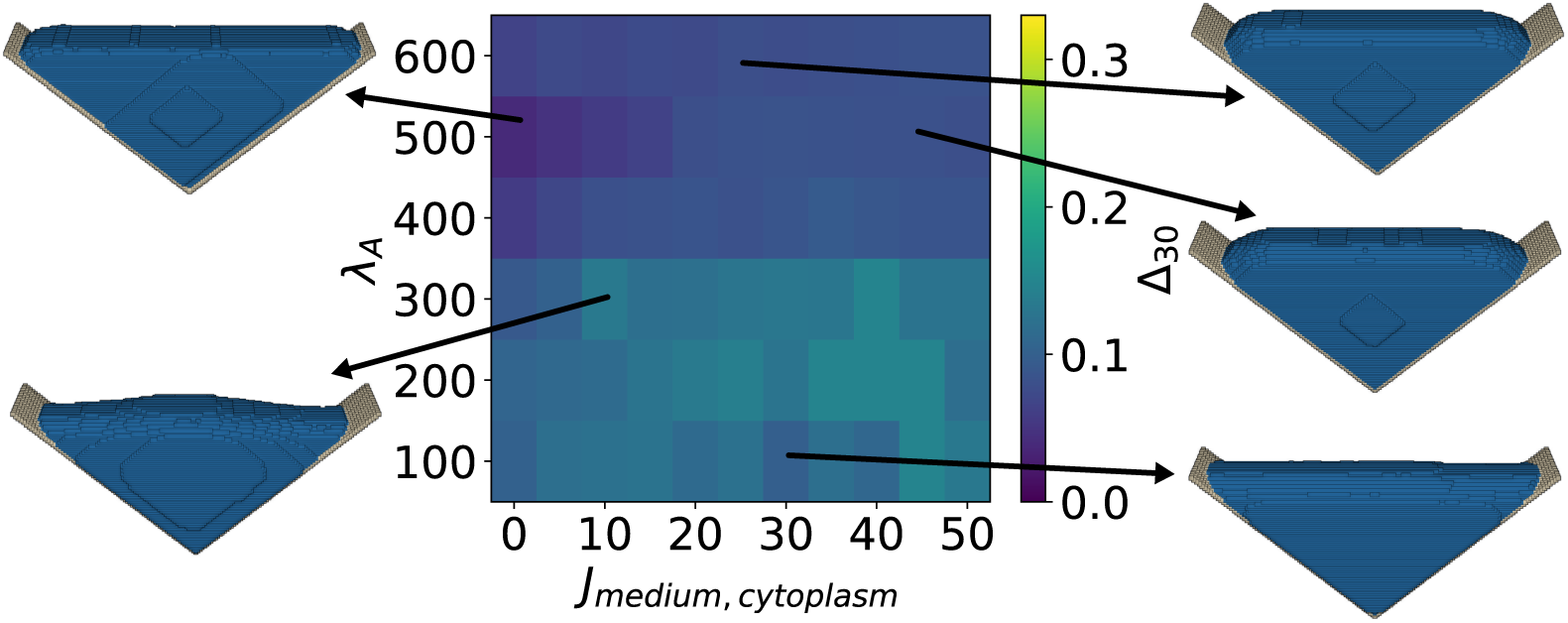
Cell shape difference Δ_30_ between the experimentally observed cell shapes and simulated cell shapes obtained with the linear area energy (Eq. 2). Differences are shown as a function of surface energy constraint *λ_A_*and interaction energy *J*_medium,cytoplasm_. Simulated cell shapes for exemplary parameter choices are presented. The minimum cell shape difference Δ_30_ = 0.039 is found for *J*_medium,cytoplasm_ = 0 and *λ_A_* = 500, the corresponding simulated cell is depicted on the top left. A movie of the simulation and final shape can be seen in S6 Video,S7 Video.

The cell shapes obtained with the elastic area Hamiltonian (see Eq. 1 and Fig. 4) are visually close to the experimentally observed cell shapes for a large parameter range. The minimum Δ_30_ = 0.090 is found for *J*_medium,cytoplasm_ = 15 and *λ_A_* = 0.05. Similar cell shapes are obtained for a large parameter range. A simulation with the elastic area Hamiltonian (Eq. 1) can be seen in S5 Video. Increasing the interaction energy *J*_medium,cytoplasm_ reduces the interface between cytoplasm and the medium, resulting in partially occupied scaffolds. For a low elastic area constraint *λ_A_*and interaction energy *J*_medium,cytoplasm_ = 0, the simulated cell shape visually differs from the experimentally observed cells and resembles again a w-shape.

The minimum found with the linear area Hamiltonian in Eq. 2 is even smaller, with Δ_30_ = 0.039 for *J*_medium,cytoplasm_ = 0 and *λ_A_* = 500, see Fig. 5 and S6 Video, S7 Video. The simulated cell shape that resembles the experiments best is triangular shaped and without invaginated arcs, however the thickness to length ratio of the cell closely resembles that of the experimentally observed shapes. Surprisingly, the simulated cell shape resembles the experimentally observed shapes best when the interaction energy *J*_medium,cytoplasm_ = 0. For a cell that is surrounded by medium only, increasing *J*_medium,cytoplasm_ has a similar effect to a linear area constraint, as the surface of the cell corresponds to the interaction area between cell and medium. In the case of structured environments, the interaction energy only acts on part of the cell surface, while the whole cell surface is relevant for the area energy. For the linear area Hamiltonian, the scaffold is always only partially covered as increasing the cytoplasm surface area always costs energy.

It is surprising that the surface corresponding best to the experiments is found for the linear area Hamiltonian simulation with interaction energy *J*_medium,cytoplasm_ = 0, as this is the simulation with the fewest parameters.

Cells with nuclei in V-shaped, right angle and equilateral triangle structures were simulated with the parameter set that performed best for the L-shaped structure (linear area Hamiltonian (Eq. 2), *λ*_A_ = 500, *J*_medium,cytoplasm_ = 0), see Fig. 6. For the right angle triangular structure, we find a good agreement between the experimentally observed shapes and the simulation with Δ_30_ = 0.053. However, for the equilateral triangle and the V-shaped structure, which differ more from the L-shaped structure than the right angle triangle, we find Δ_30_ = 0.141 and Δ_30_ = 0.136. These differences show that it is difficult to find universal parameters for the CPM that accurately predict cell shape in different structured environments.

**Fig 6.**
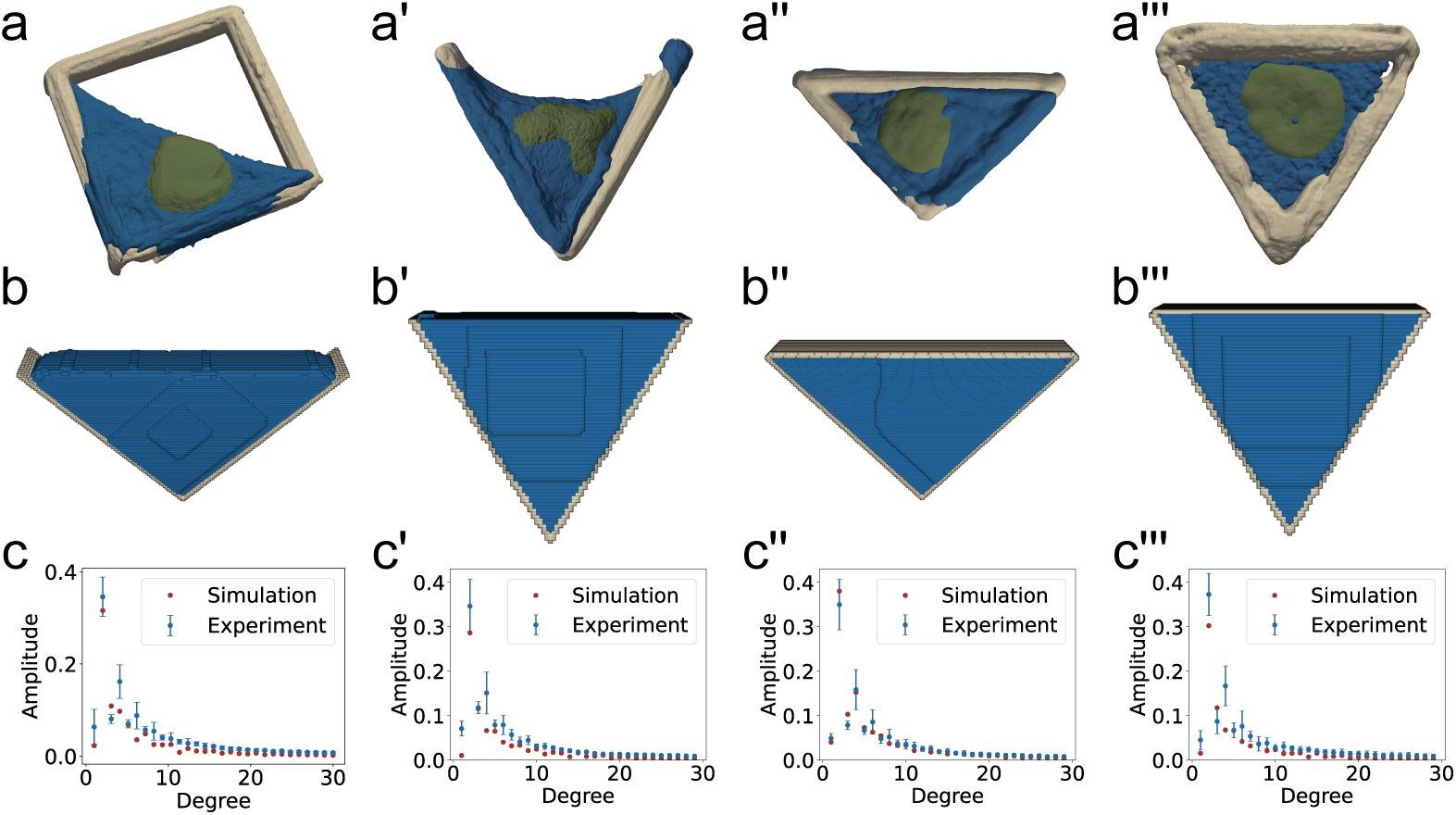
Experimental and simulated shapes and frequency spectrum. **a-a***^′′′^*) Reconstruction of the experimental data from cells in L-shaped, V-shaped, right-angle triangular and equilateral triangular shaped structures. **b-b***^′′′^*) Simulated cell shapes in structured environments. **c-c***^′′′^*) Frequency spectra of the experimental and simulated cell shapes. **c)** Δ_30_ = 0.039, **c***^′^*) Δ_30_ = 0.136, **c***^′′^*) Δ_30_ = 0.053 **c***^′′′^*) Δ_30_ = 0.141.

From the experimental data, we find that the nucleus is significantly deformed by the cell cytoskeleton, but does not have a large effect on cell shape. We include the nucleus explicitly in our CPM simulations, and do not find a visual effect of the nucleus on cell shape in the simulations. In S1, we simulate cells without explicit nucleus representation in the CPM, and find only minor cell shape changes.

## Discussion

Due to recent technical advances, an increasing amount of experimental data is three-dimensional (3D). This adds to the need for 3D single cell models to explain and predict cell shape. In this work, we compared different approaches to describe cell shape in 3D. We used surface minimization and different variants of the cellular Potts model (CPM) to investigate how much simulated shape predictions differ from experimentally observed cell shapes in precisely defined environments. We find that the predicted constant mean curvature (CMC) shapes agree well if the cell does not form invaginated arcs and stress fibers. Larger differences were found for cell shapes in L- and V-shaped structures.

Better cell shape prediction was possible with the CPM, independent of the choice of Hamiltonian and specific parameter values. Invaginated arcs form without explicitly simulating stress fibers and the high quadrupole moment in the spherical harmonics is correctly predicted. There are multiple factors that contribute to the better performance of CPM simulations compared to surface minimization models. By selecting the neighbor order in the CPM, one can define how local the interaction energy is. This influences to which the surface minimization can occur, or may even facilitate surface extension between two general cell types, if the interaction energy is negative. A cell described by a Hamiltonian with volume, area, interaction and nucleus centering energies leads to more complex shapes that better resemble experimental results.

Even though CPM simulations better capture experimental cell shapes, it should be noted that there are some systematic differences between the simulated and experimental shapes. For instance, cells in experiments have more variability in their shapes and attach only partially to the structures compared to the smooth shapes predicted by the CPM. Additionally, the reconstruction of cell shape in experiments using actin staining is only an approximation, and may introduce some degree of error in the obtained cell shapes. Actin stress fibers often span between adhesive sites, however their impact has not been accounted for in the simulations and could be included in the future.

The best agreement between experimental and simulated cell shape was found with the linear area Hamiltonian, despite the fact that less parameters are used during the simulation. The effect of the nucleus on cell shape is neglectable in this setting, as the nucleus is strongly deformed by the cytoskeleton. In the future, these results could be used for rational scaffold design which then can be implemented with additive manufacturing methods like the 3D laser nanoprinting used here.

## Supporting information

**S1 Appendix. Cell shapes simulated with the CPM without explicit representation of the nucleus.**

**S1 Table. Parameters for the CPM simulations.**

**S1 Video. Shape of an experimentally observed single cell in an L-shaped structure (left) and minimized surface in an L-shaped structure (right).**

**S2 Video. Shape of an experimentally observed single cell in a V-shaped structure (left) and minimized surface in a V-shaped structure (right).**

**S3 Video. Shape of an experimentally observed single cell in a right-angle triangular structure (left) and minimized surface in a right-angle triangular structure (right).**

**S4 Video. Shape of an experimentally observed single cell in an equilateral triangular structure (left) and minimized surface in an equilateral triangular structure (right).**

**S5 Video. Cellular Potts model simulation of a single cell in an L-shaped structure with the elastic area Hamiltonian (Eq. 1).**

**S6 Video. Cellular Potts model simulation of a single cell in an L-shaped structure with the linear area Hamiltonian (Eq. 2).**

**S7 Video. Result of the cellular Potts model simulation of a single cell in an L-shaped structure with the linear area Hamiltonian (Eq. 2).**

## Code and data availability

CompuCell3D simulation scripts and the linear surface plugin for clusters as well as python scripts to generate input files for SurfaceEvolver from STL-files and to obtain the frequency spectrum from cell shapes will be available from zenodo.org upon publication.

## Acknowledgments

This work is supported by the Deutsche Forschungsgemeinschaft (DFG, German Research Foundation) under Germany’s Excellence Strategy EXC 2082/1-390761711 (cluster of excellence 3DMM2O). USS is a member of the Interdisciplinary Center for Scientific Computing (IWR).

## Competing interests

The authors declare no competing interests.

## Author Contributions

**Conceptualization:** Rabea Link, Mona Jaggy, Martin Bastmeyer, Ulrich S. Schwarz.

**Data curation:** Rabea Link, Mona Jaggy.

**Formal analysis:** Rabea Link.

**Funding acquisition:** Martin Bastmeyer, Ulrich S. Schwarz.

**Investigation:** Rabea Link, Mona Jaggy.

**Methodology:** Rabea Link, Mona Jaggy.

**Project administration:** Martin Bastmeyer, Ulrich S. Schwarz.

**Resources:** Martin Bastmeyer, Ulrich S. Schwarz.

**Software:** Rabea Link.

**Supervision:** Martin Bastmeyer, Ulrich S. Schwarz.

**Validation:** Rabea Link, Mona Jaggy.

**Visualization:** Rabea Link, Mona Jaggy.

**Writing – original draft:** Rabea Link, Ulrich S. Schwarz.

**Writing – review & editing:** Rabea Link, Mona Jaggy, Martin Bastmeyer, Ulrich S. Schwarz.

## Notes

### Competing Interest Statement

The authors have declared no competing interest.

